# The parasitic plant *Cuscuta campestris* selectively loads *trans*-species miRNAs onto host Argonautes, but not self Argonautes

**DOI:** 10.1101/2025.10.16.682822

**Authors:** Ya-Chi Nien, Michael J. Axtell

## Abstract

The obligate parasitic plant *Cuscuta campestris* delivers *trans*-species microRNAs (miRNAs) into host plants to silence host mRNAs. Here, the genetic requirements for biogenesis, movement, and function of these miRNAs were investigated. Primary miRNA transcript accumulation precedes mature miRNA accumulation by 24 to 48 hours. *Trans-*species miRNAs accumulate in host tissues a short distance from the site of parasite attachment. *Trans-*species miRNAs require *C. campestris* but not host *Dicer-Like 1* (*DCL1*) for accumulation. These miRNAs specifically avoid Argonaute (AGO) loading in *C. campestris* tissue where they instead accumulate as miRNA/miRNA* duplexes. After arrival and short-distance spreading in host tissues, they are selectively loaded into host AGO1. This study clarifies the transcription, dicing, delivery, and function of *C. campestris trans*-species miRNAs. We propose that selective avoidance of self AGO-loading is a mechanism to facilitate the delivery of these “export only” miRNAs to host tissues.

## Introduction

*Cuscuta* is a genus of parasitic plants with minimal chlorophyll and no functional leaves or roots, producing yellow to orange vines. As obligate holoparasites, they rely on host plants to complete their life cycle^1^. The *Cuscuta* genus contains approximately 200 species with a diverse host range^2^. Some species exhibit narrow host preferences, while others, like *Cuscuta campestris*, parasitize a broad spectrum of plants—including most eudicots and some monocots^3^. Its broad host range makes *C. campestris* an agricultural nuisance that can cause substantial crop losses. Understanding the virulence mechanisms of *Cuscuta* is therefore critical for developing effective crop protection strategies.

The haustorium is a specialized organ that anchors *Cuscuta* to its host and primarily functions as a vascular connection for the withdrawal of water, photosynthate, and nutrients from the host plant. The haustorium also serves as a conduit for extensive molecular exchange including RNAs^4^. Previous studies have shown that microRNAs (miRNAs) can also move from the parasite into host tissues where they can regulate host mRNAs^5–8^.

Canonical plant miRNAs are usually 21 or 22 nucleotides in length and are processed from single-stranded precursors that fold into characteristic hairpin structures. These hairpins are processed by the endonuclease Dicer-Like 1 (DCL1) to produce miRNA/miRNA* duplexes. The mature miRNA is loaded onto an Argonaute (AGO) protein, often AGO1, to form the RNA-induced silencing complex (RISC), while the miRNA* is subsequently degraded^9^. The RISC mediates sequence-specific cleavage and/or translational repression of target mRNAs, leading to gene silencing at the post-transcriptional level^9,10^. Gene regulation by canonical plant miRNAs collectively impacts many biological functions, including development^11^, organogenesis, inorganic nutrition^12^, and pathogen defense^13^. Other categories of short interfering RNAs (siRNAs) can also be loaded onto AGO proteins to specify gene silencing. Plant siRNAs are often produced from double stranded RNA precursors by one or more alternative DCL proteins (DCL2, 3, or 4).

Exchange of functional miRNAs and siRNAs between plants and their associated organisms has been reported in diverse systems, including plant-plant, plant–fungus, plant–insect, and plant–mammal interactions^5,14^. One well-studied example is the fungal grey mold pathogen *Botrytis cinerea*, which delivers siRNAs into host plants to suppress immunity^6^, while plants counter by exporting siRNAs and miRNAs into the fungus to reduce virulence^7^. In both cases, the mobile small RNAs are processed in the donor organism—requiring DCL1/2 in the pathogen and DCL2/3/4 in the host plant. The mobile small RNAs travel within extracellular vesicles (EVs) and incorporate into an AGO protein upon arrival^7,8^. However, the nature of the mobile agent may differ between the two organisms. Exported *A. thaliana*-derived small RNAs require RNA-binding proteins, such as AGO1, RNA helicases, and annexins, to be sorted into and stabilized within EVs^15^. These proteins are bound to the small RNAs and co-localize with the EVs, suggesting that plant sRNAs are transported to *B. cinerea* as sRNA-protein complexes. In contrast, for fungal-derived siRNAs, no direct evidence describes their transport form, but successful incorporation into host AGO1 suggest they may move as free duplexes.

*C. campestris* expresses a cohort of 96 distinct miRNA families at high levels specifically in the haustorium^16–18^. All of the *MIRNA* genes for these interface-induced miRNAs have a common promoter element which is identical to that used to drive plant small nuclear RNA (snRNA) transcription^18^. This promoter element drives transcription by RNA polymerase III which marks the *C. campestris* interface-induced *MIRNA* genes as distinct from canonical plant *MIRNA* genes: Canonical *MIRNA* genes in plants are transcribed by RNA polymerase II^19^. At least a subset of the interface-induced miRNAs accumulate in the host where they target host transcripts for cleavage and secondary siRNA production, potentially modulating host gene expression to facilitate parasitism^17^. We refer to these as *trans*-species miRNAs. Despite extensive efforts, we have found no evidence that *C. campestris trans*-species miRNAs are functional within the parasite^17,20^, suggesting they are used for “export only”.

Here we investigate the biogenesis and genetic requirements for *C. campestris trans*-species miRNA function. Rapid accumulation of *trans-*species miRNAs in haustoria is preceded by accumulation of primary transcripts. Mature *trans-*species miRNAs accumulate a short distance (∼1-2 centimeters) in host tissues adjacent to the haustorium. Parasite *DCL1*, but not host *DCL1*, is required for accumulation of *trans*-species miRNAs. Once made, *trans-*species miRNAs are specifically prevented from associating with parasite AGO proteins and instead accumulate as free miRNA/miRNA* duplexes inside of the parasite. Upon arrival in the host, they are loaded into host AGO1. The selective avoidance of self-AGO loading may be a mechanism to promote export of *C. campestris trans-*species miRNAs.

## Results

### *Trans*-species miRNA accumulation extends into adjacent host stem

*Cuscuta campestris* accumulates *trans*-species miRNAs specifically at the haustorial interface^17^, some of which are detectable in host tissues many centimeters away^21^. To gain quantitative estimates of *trans-*species miRNA accumulation in various tissues, two representative *C. campestris trans*-species miRNAs: ccm-miR12480b, which targets *SEOR1*, and ccm-miR12497f, which targets *TIR1, AFB2*, and *AFB3*^17,20^ were examined. As expected, both miRNAs accumulated most strongly at the haustorial interface (**Fig. 1**). Notably, substantial amounts were also detected in adjacent host petioles, both proximal and distal to haustoria, at ∼10-fold lower levels than at the interface (**Fig. 1**). Accumulation drops sharply (∼10,000-fold lower) at the petiole base, indicating locally limited movement of *C. campestris*-derived *trans*-species miRNAs into host tissue.

**Figure 1.**
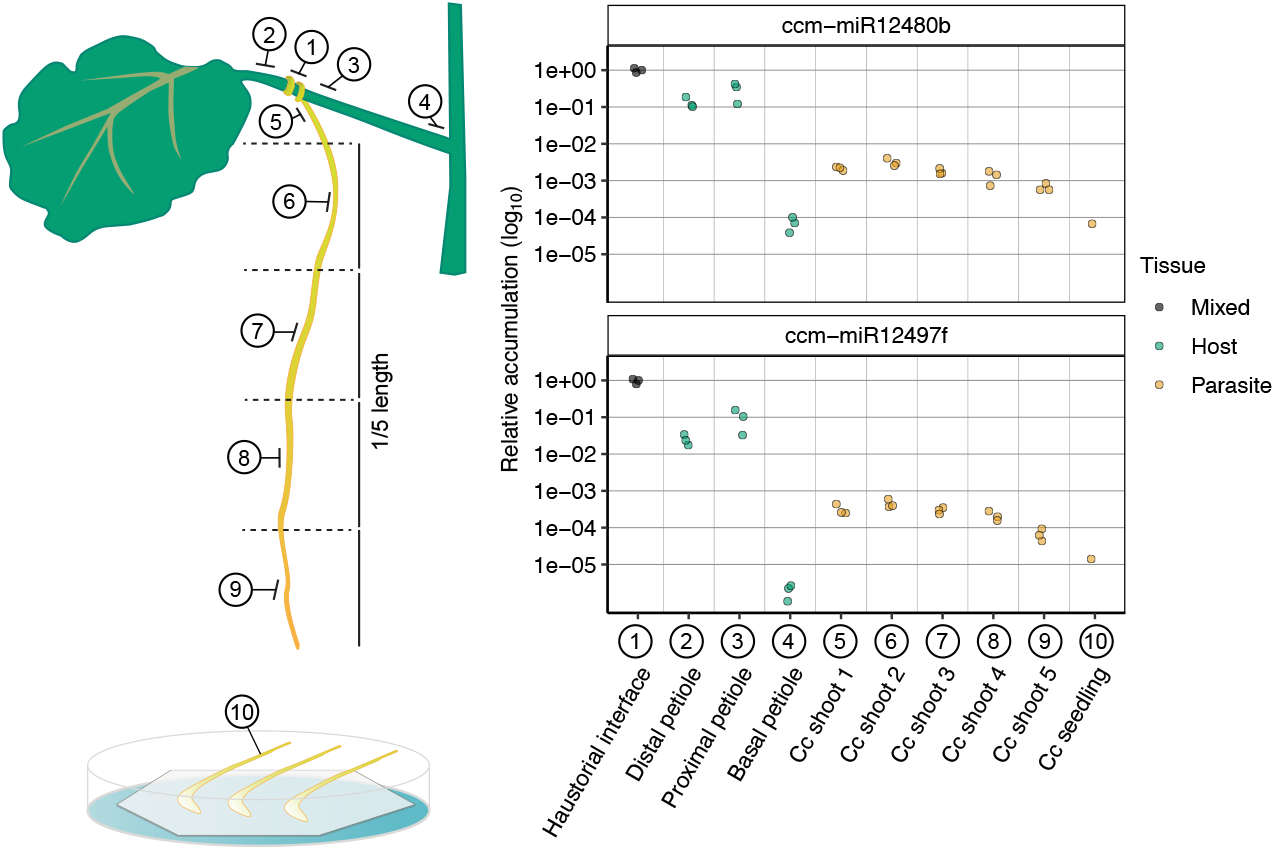
*Trans*-species miRNAs accumulate a short distance from the haustorial attachment. RT-qPCR quantified mature *trans*-species miRNAs in: (1) 10-day-old interface tissue containing *C. campestris* haustoria and *N. benthamiana* petiole beneath the attachment; (2–3) petiole segments located 1 cm distal or proximal to the attachment; (4) basal petiole adjacent to the *N. benthamiana* stem; (5–9) five 1 cm *C. campestris* shoot segments extending from the attachment to parasite tips; and (10) 2.5-day-old *C. campestris* seedlings. Colors distinguish tissue origin: gray, mixed host–parasite tissue; green, *N. benthamiana* only; yellow, *C. campestris* only. Each dot represents a biological replicate (three per condition, except for *C. campestris* seedling, which has one sample). Relative accumulation was normalized to haustorial interface by the 2^−ΔCt^ method. Data available in **Supplementary Table 1**.

Within the parasite, *trans*-species miRNAs were detectable throughout the *C. campestris* shoot, at ∼5,000-fold lower levels than at the interface and declining further toward the shoot tips. Minimal accumulation was observed in seedlings (**Fig. 1**). These results reveal that accumulation extends beyond the haustorial interface into non-haustorial parasite and host tissues.

### Primary transcripts peak 24–48 hours before mature miRNAs and do not accumulate in host tissues

Mature *C. campestris trans*-species miRNAs accumulate in haustoria 72–96 hours after the onset of haustoria development, showing temporal specificity and accumulating regardless of host presence^18^. They reach ∼10-fold higher levels than adjacent host tissue and ∼5000-fold higher than *C. campestris* shoots (**Fig. 1**), highlighting spatial enrichment. We investigated the mechanism controlling this precise temporal and spatial pattern. Four factors together determine the net accumulation of mature miRNA: (1) transcription rate of the primary transcript, (2) non-dicing degradation of the primary transcript, (3) dicing rate which processes primary transcripts into mature miRNAs, and (4) the mature miRNA degradation rate. It’s difficult to determine which factors drive net accumulation, and the observed pattern may result from a combination of all four. As a first step, we measured the levels of primary transcript across tissues and developmental stages that vary in mature miRNA abundance (**Fig. 2A, B**).

**Figure 2.**
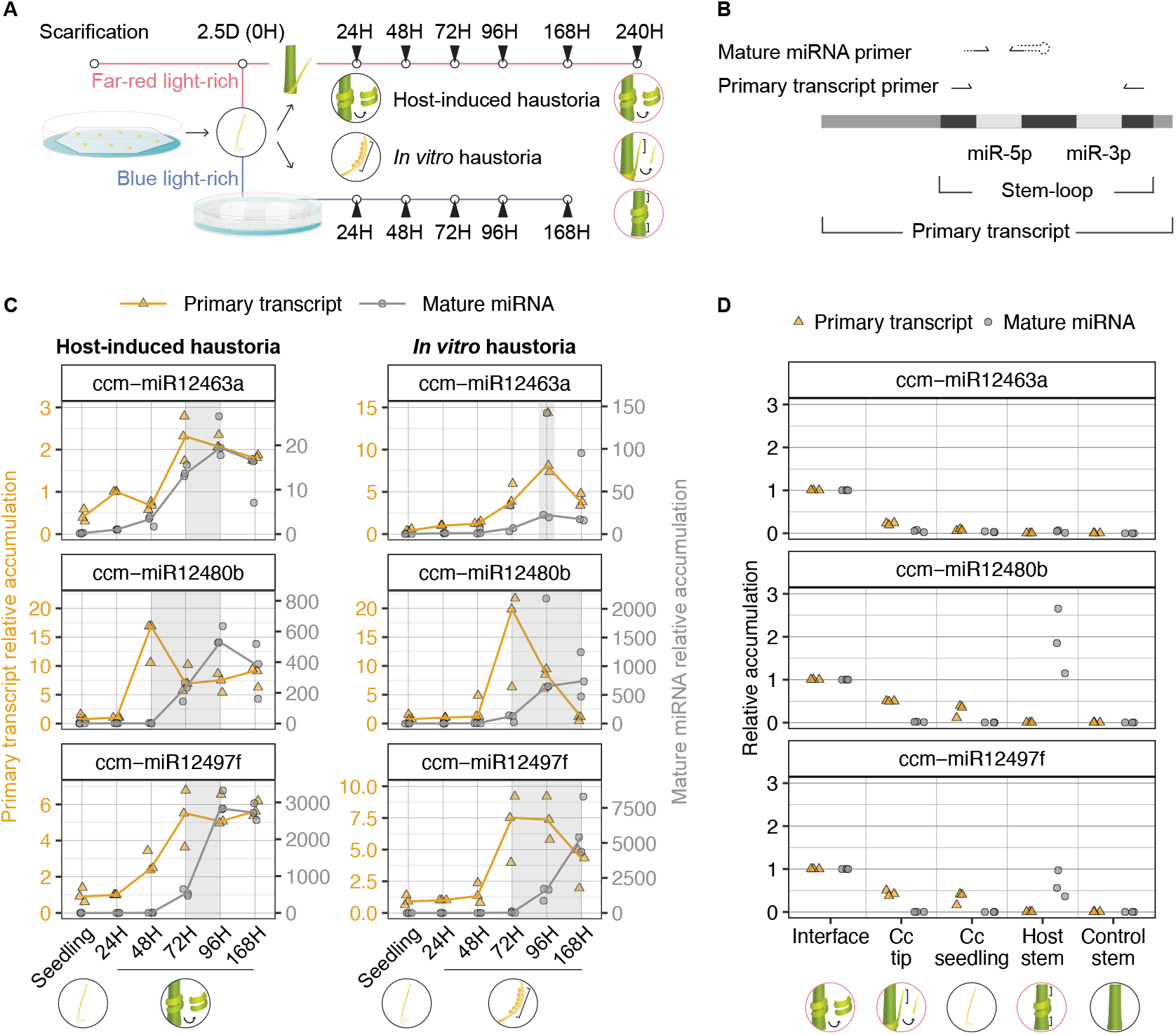
Temporal and spatial accumulation of *trans*-species miRNAs and their primary transcripts. **(A)** Experimental design. *C. campestris* seedlings were either attached to *A. thaliana* under far-red light enriched conditions (host-induced haustoria, pink line) or sandwiched with microscope slides and agar under blue light (*in vitro* haustoria, blue line). *C. campestris* seedling and haustoria from both treatments were collected at 24, 48, 72, 96, and 168 hours (black-bordered circles) and 240 hours (pink-bordered circles) post-induction. **(B)** RT-qPCR primer design for detecting mature miRNAs and primary transcripts. Solid lines indicate exact-match nucleotides; dashed lines indicate universal or random nucleotides added to achieve the desired melting temperature. **(C)** Temporal profiles of primary transcript (yellow) and their mature miRNA (gray) in host-induced haustoria (left) and *in vitro* haustoria (right). Shaded boxes indicate the time lag between primary transcripts and mature miRNA peak expression. Each dot is a biological replicate consisting of 5-20 pooled specimens, with three replicates per condition. **(D)** Spatial analysis of primary transcript (yellow) and their mature miRNA (gray). *C. campestris* uncoiled stems at the interface, *C. campestris* cut tips, and adjacent host stems, were dissected from 240-hour host-induced haustorium, alongside non-parasitized *C. campestris* seedlings and *A. thaliana* control stems. Each dot is a biological replicate consisting of 5-20 pooled specimens, with three replicates per tissue. Expression levels in (C) and (D) were calculated using the 2^-ΔΔCt^ method and 2^-ΔCt^, respectively, with *C. campestris* snRNA U6 as the reference gene for primary transcripts and miR159 for mature miRNAs; the 24-hour sample and haustorial interface were used as calibrators, respectively. Data available in **Supplementary Table 1**.

In addition to the previously measured miRNAs (ccm-miR12480b and ccm-miR12497f), an additional *trans*-species miRNA, ccm-miR12463a, which targets *BIK1*^16,17^, was examined. We found that primary transcripts, in both host-induced haustoria and host-free *in vitro* haustoria, generally peak 24-48 hours before the highest accumulation of mature miRNAs (**Fig. 2C**). This temporal lag suggests that primary transcripts accumulate before mature miRNAs appear. Since the lag occurs in both haustoria types, the control mechanism is likely intrinsic to the parasite’s developmental program rather than dependent on host cues (**Fig. 2C**). Spatially, primary transcript levels are highest at the haustorial interface, matching the peak of mature *trans*-species miRNA (**Fig. 2D**). In *C. campestris* tips and seedlings, where mature miRNAs were low or undetectable, primary transcripts are also relatively low but still detectable at 25-50% of the interface level (**Fig. 2D**). This suggests that in tissues with the potential to form haustoria, such as seedlings and shoot tips, a basal level of primary transcripts is produced and poised for processing into mature miRNA upon haustoria onset.

Although primary transcripts and mature miRNAs are generally positively correlated, an exception occurs in the host stem adjacent to the haustoria, where only mature miRNAs—but not primary transcripts—were detected (**Fig. 2D**). This pattern was observed for two of the three *trans*-species miRNAs tested. The exception is ccm-miR12463a, the lowest-expressed of three miRNAs examined. Mature *trans*-species miRNAs spread into adjacent host tissue at roughly 10-fold lower level (**Fig. 1**). Because ccm-miR12463a is already expressed at ∼20- and 200-fold lower level than the other two miRNAs in the host-induced haustoria (**Fig. 2C**), its translocated miRNA in the adjacent host stem may have fallen below the detection threshold. Taken together these data suggest that either duplex miRNA/miRNA* or single-stranded mature miRNAs, rather than their precursors, move into the host. These results indicate that *trans*-species miRNAs are both transcribed and processed within the parasite using its own machinery.

### Parasite *DCL1*, but not host *DCL1*, is required for *trans*-species miRNA accumulation

To further investigate the contribution of parasite versus host machinery in *trans*-species miRNA production, we targeted a key miRNA biogenesis factor, *DCL1*, in either the parasite or the host. A reduction of mature miRNAs upon parasite *DCL1* knockdown would indicate processing within the parasite, whereas a decrease following host *DCL1* silencing would suggest that primary transcripts are translocated and processed in the host. Previous studies have shown that artificial miRNA (amiR) can move from agroinfiltrated leaves to non-infiltrated leaves, silencing genes in both leaves, likely via the vascular continuum of the petiole and stem^22,23^. Recognizing the potential of this mobility, we built on the concept of host-induced gene silencing by using *Agrobacterium tumefaciens* infiltration (“agroinfiltration”) into *Nicotiana benthamiana* host leaves. Agroinfiltration with strains encoding amiR was predicted to result in systemic amiR migration into adjacent petioles. Attachment of *C. campestris* to such petioles allows amiR delivery to the parasite (**Fig. 3**).

**Figure 3.**
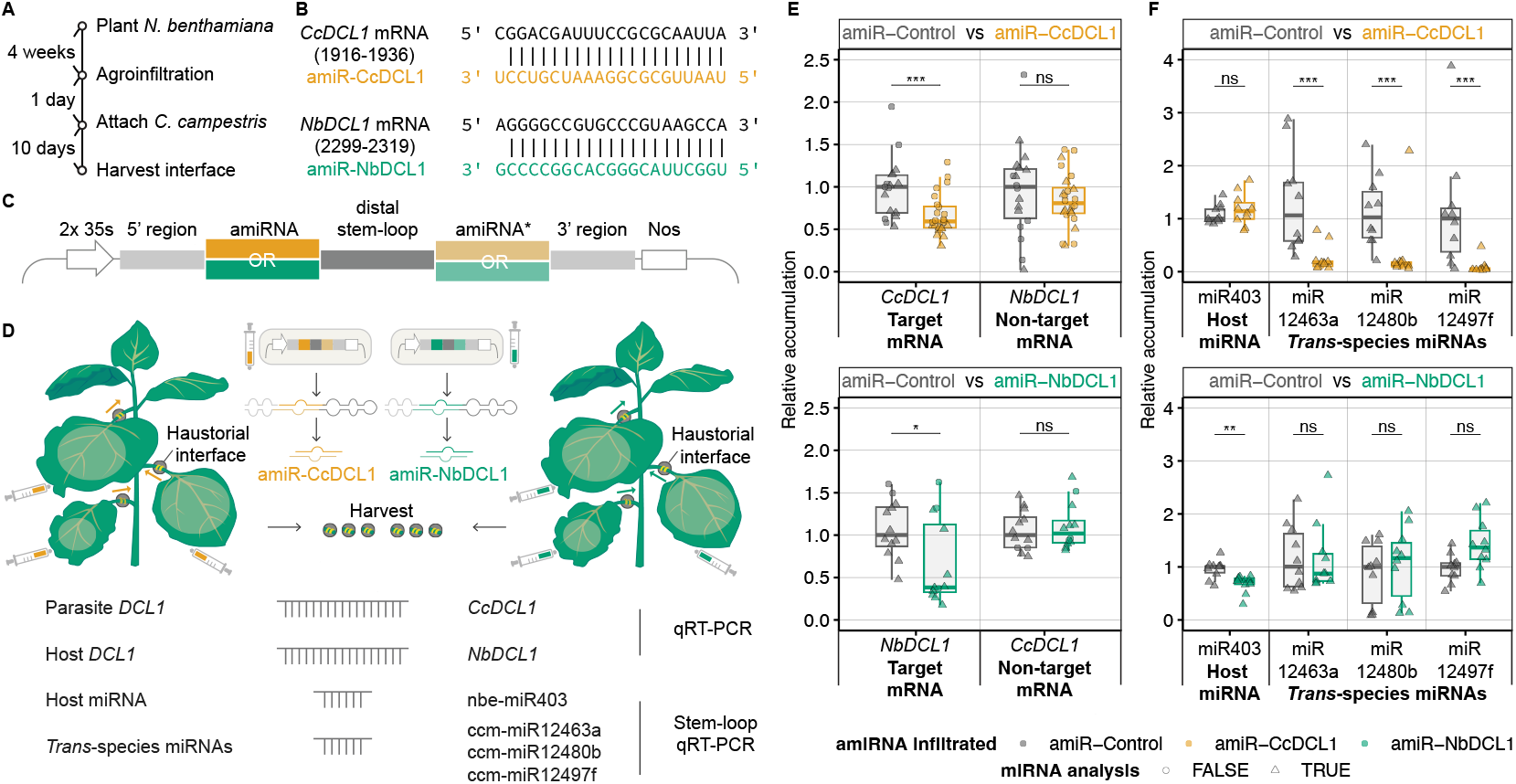
Parasite *DCL1*, but not host *DCL1*, is required for *trans*-species miRNA biogenesis. **(A)** Workflow for artificial miRNA (amiR)-mediated gene silencing. **(B)** Alignment of amiRs with their targets. **(C)** Schematic of the amiR vector. **(D)** Application of amiR and subsequent RT-qPCR analysis. **(E)** Expression of *CcDCL1, NbDCL1*, **(F)** a canonical host miRNA, and three *trans*-species miRNAs following infiltration with control amiR (gray), *CcDCL1*-targeting amiR (yellow), or *NbDCL1*-targeting amiR (green). Each dot is a biological replicate made from 3-6 pooled specimens. Expression values were calculated using the 2^−ΔΔCt^ method. Reference gene was *CcRPN7*^24^ and *NbPP2A*^25^ for mRNA analysis and Cc 5.8s rRNA and Nb 5.8s rRNA for miRNA analysis. The amiR-Control samples were used as calibrators. One-tailed Mann–Whitney U test was used for pairwise comparisons. Statistical significance is indicated: *** p < 0.001, ** p < 0.01, * p < 0.05, ns not significant. Data available in **Supplementary Table 1**.

Having observed that both *C. campestris* and *N. benthamiana* encode a single copy of *DCL1* (**Extended Data Fig. 1**), we designed species-specific artificial miRNAs targeting parasite (amiR-CcDCL1) and host *DCL1* (amiR-NbDCL1) (**Fig. 3B**) and cloned them into an *A. thaliana MIR390a* backbone (**Fig. 3C**). The amiR construct was agroinfiltrated into *N. benthamiana*, followed by *C. campestris* attachment to the petiole of the infiltrated leaf, and haustorial tissues were collected after 10 days (**Fig. 3A, D**). Targeting *DCL1* with species-specific amiR knocked down parasite and host *DCL1* by 40% and 50%, respectively, while non-targeted *DCL1* was unaffected, indicating silencing without detectable off-target effects (**Fig. 3E**). *C. campestris* establishment rates were lower on amiR-CcDCL1 *N. benthamiana* and this effect was statistically significant in two of five trials (**Extended Data Fig. 2**). This suggested that the strongest *CcDCL1* silencing may impact *C. campestris* establishment. Our goal was to evaluate the impact of *trans*-species miRNAs under conditions where *CcDCL1* and *NbDCL1* were effectively suppressed. To achieve this, we analyzed the top ten living specimens with the strongest knockdown of *CcDCL1* and *NbDCL1* for miRNA quantification. As expected, a host canonical miRNA that is not encoded by the parasite genome, nbe-miR403, was dependent on *NbDCL1* (**Fig. 3F**). The three tested *trans*-species miRNAs decreased significantly with parasite *DCL1* knockdown but remained unchanged when host *DCL1* was reduced, indicating that parasite *DCL1* is required for their biogenesis, whereas host *DCL1* is dispensable (**Fig. 3F**).

### *C. campestris trans*-species miRNAs accumulate as miRNA/miRNA* duplexes within parasite tissue

Since mature *trans*-species miRNA are produced within the parasite, the key question is whether the mobile form that crosses the species barrier is single-stranded, duplex miRNA/miRNA*, and/or AGO-bound. To test this, we examined the ratio of miRNA to miRNA* present in the parasite prior to export. A higher proportion of the miRNA, as seen for most canonical miRNAs, would suggest movement as a single-stranded or AGO-bound form. In contrast, an approximately equal (∼50%) distribution of miRNA and miRNA* would implicate a duplex as the mobile agent. Three replicate small RNA-seq libraries from *C. campestris* host-free *in vitro* haustoria were previously shown to accumulate high levels of *trans*-species miRNAs^18^. These data were analyzed by determining mature miRNA and miRNA* accumulation for each miRNA family and calculating the percentage of mature miRNA relative to the total reads (miRNA + miRNA*) **(Fig. 4A)**. This percentage was significantly higher for canonical miRNAs compared to *trans*-species miRNAs **(Fig. 4B)**. The median value for *trans*-species miRNA families was around 50% of mature miRNA, with a large variance. The typical canonical miRNA family, in contrast, had more than 90% of the reads as mature miRNA with much less miRNA* **(Fig. 4B)**. These findings suggest that *trans*-species miRNAs predominantly exist as miRNA/miRNA* duplexes within the parasite, raising the possibility that they are not loaded onto parasite AGO proteins and are instead translocated across the host–parasite interface in duplex form.

**Figure 4.**
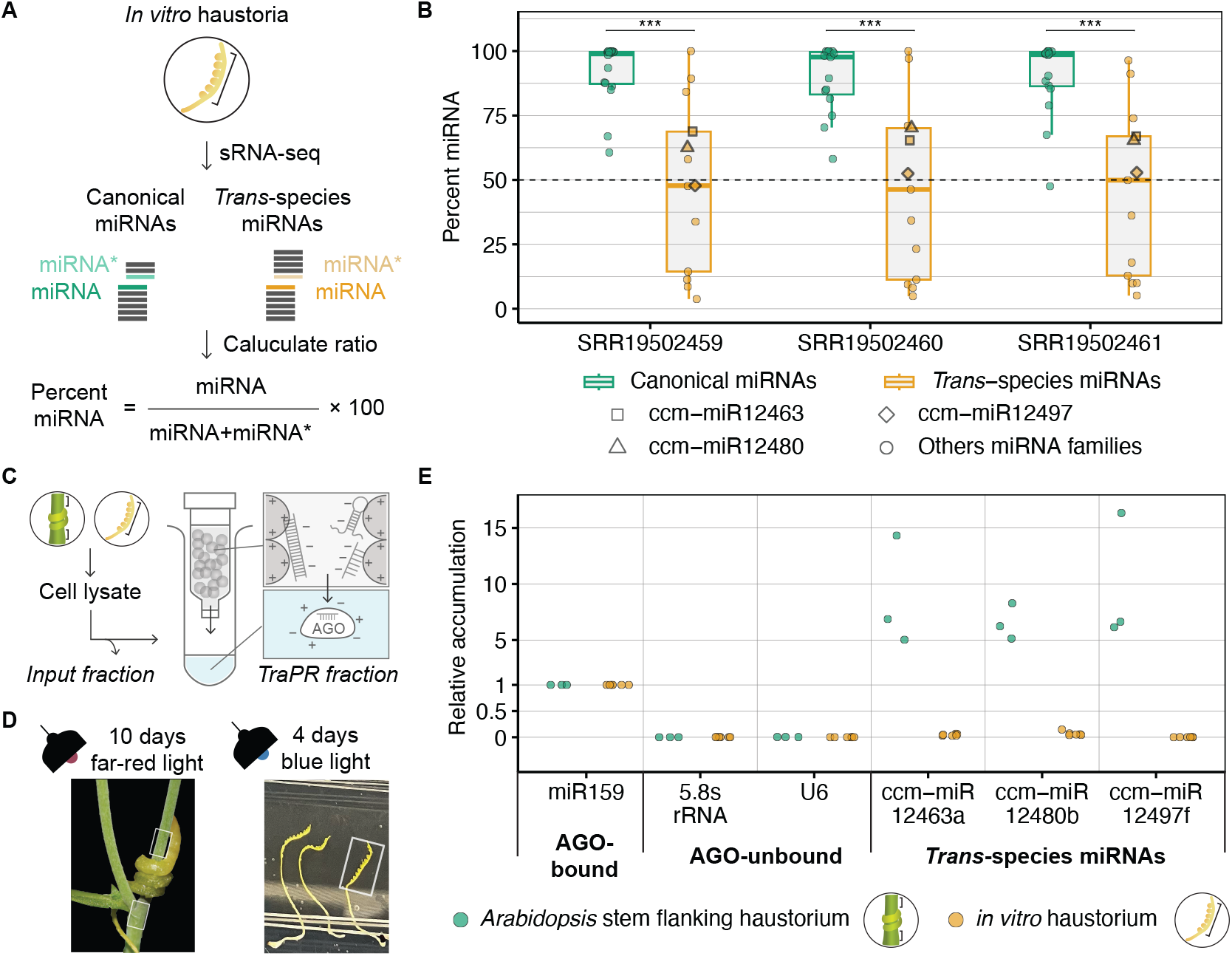
Strand ratio and AGO association of *trans*-species miRNAs in *C. campestris* and host tissues. **(A)** Workflow to calculate percent mature miRNA from sRNA-seq libraries from host-free *C. campestris in vitro* haustoria. **(B)** Percent miRNA in three sRNA-seq libraries. Each dot represents a miRNA family. The three *trans*-species miRNA families analyzed elsewhere in this study are highlighted: square = ccm-miR12463, triangle = ccm-miR12480, diamond = ccm-miR12497. The dashed line marks 50%. Mann-Whitney test was used for pairwise comparisons. Statistical significance is indicated: *** p < 0.001. **(C)** TraPR workflow. A portion of total RNA is kept as input fraction. The remaining RNA is incubated with positively charged resin to trap unprotected RNAs. Protein-RNA complexes escape the resin and elute into TraPR fraction. **(D)** Tissue collection of host-induced haustoria (left) and host-free *in vitro* haustoria (right). Harvested regions are boxed. **(E)** RT-qPCR analysis of RNAs from host stem adjacent to haustoria (green) and *in vitro* haustoria (yellow). Each dot in host stem samples represents pooled RNA from ∼36 *A. thaliana* stem segments, half from above and half from below the haustorial attachment. Each dot in the *in vitro* haustoria samples represent pooled RNA from 60–70 *C. campestris* seedlings. Each condition included 3–6 biological replicates. Relative expression was calculated using the 2^−ΔΔCt^ (TraPR – Input fraction) method, with miR159 as housekeeping gene. Data available in **Supplementary Table 1**.

### *Trans*-species miRNAs are not AGO-bound in the parasite but are AGO-bound in host tissues

The approximately equal accumulation of miRNA and miRNA* suggests that *trans*-species miRNAs exist as miRNA/miRNA* duplexes within the parasite. One mechanism to explain this would be avoidance of loading onto *C. campestris* AGOs. To test this hypothesis, we used the TraPR (**T**rans-kingdom, **r**apid, **a**ffordable **P**urification of **R**ISCs) method. This method relies on positively charged resin that captures free, unprotected RNAs, while protein-bound RNAs pass through and become enriched (**Fig. 4C**). Two tissue types were analyzed (**Fig. 4D**). First, *A. thaliana* host stem adjacent to *C. campestris* attachment sites, where the spread of *trans*-species miRNA was confirmed and shown to reach detectable levels. Second, host-free *in vitro* haustoria induced by blue light for 96 hours were examined. At this stage, *trans*-species miRNAs are already produced at detectable levels comparable to those in host-induced haustoria (**Fig. 2C**). These samples contain only *C. campestris* tissue, derived from seedlings that never encounter a host, providing a naïve environment where only parasite AGOs can interact with the *trans*-species miRNAs.

In both tissues, the known AGO1-bound miR159 was enriched, while unbound RNAs such as 5.8s rRNA and U6 snRNA were depleted (**Fig. 4E**), confirming that TraPR distinguishes bound from unbound RNAs. The three tested *trans*-species miRNAs were strongly enriched in the host stem but depleted in *in vitro* haustoria (**Fig. 4E**). These results indicate that *trans*-species miRNAs are not bound to AGO in the parasite-only environment but become AGO-bound after export into host. We conclude that *C. campestris trans*-species miRNAs are selectively loaded into host AGOs, but not into parasite AGOs.

### *C. campestris trans*-species miRNAs are preferentially loaded into host AGO1

After establishing that *C. campestris trans*-species miRNAs are AGO-bound only in host tissues, the next step was to determine which host AGO loads them. To test this, host AGOs were enriched by immunoprecipitation, and associated RNAs were examined. Ccm-miR12463a, ccm-miR12480b, and ccm-miR12497f have all been shown to direct slicing of *A. thaliana* mRNAs^17,20^, and each has a 5’-U. These features are consistent with AGO1 loading. AGO2 was also of interest because of its role in plant defense^26,27^ against viruses and bacteria. *C. campestris* was attached to *A. thaliana* lines expressing epitope-tagged AGO1 or AGO2, as well as to wild-type plants. The parasite was then removed after ten days, and AGO immunoprecipitation was performed using specimens collected from the host stem beneath the attachment site (**Fig. 5A**). AGO1-bound ath-miR166 and AGO2-bound ath-miR390 / ath-miR408 were enriched in their respective lines, confirming assay robustness (**Fig. 5B**). All three *C. campestris trans*-species miRNAs were strongly enriched in AGO1 pulldowns, with much weaker signals in AGO2 pulldowns (**Fig. 5B**). These results indicate that these three *C. campestris trans*-species miRNA preferentially loaded on host AGO1 over AGO2.

**Figure 5.**
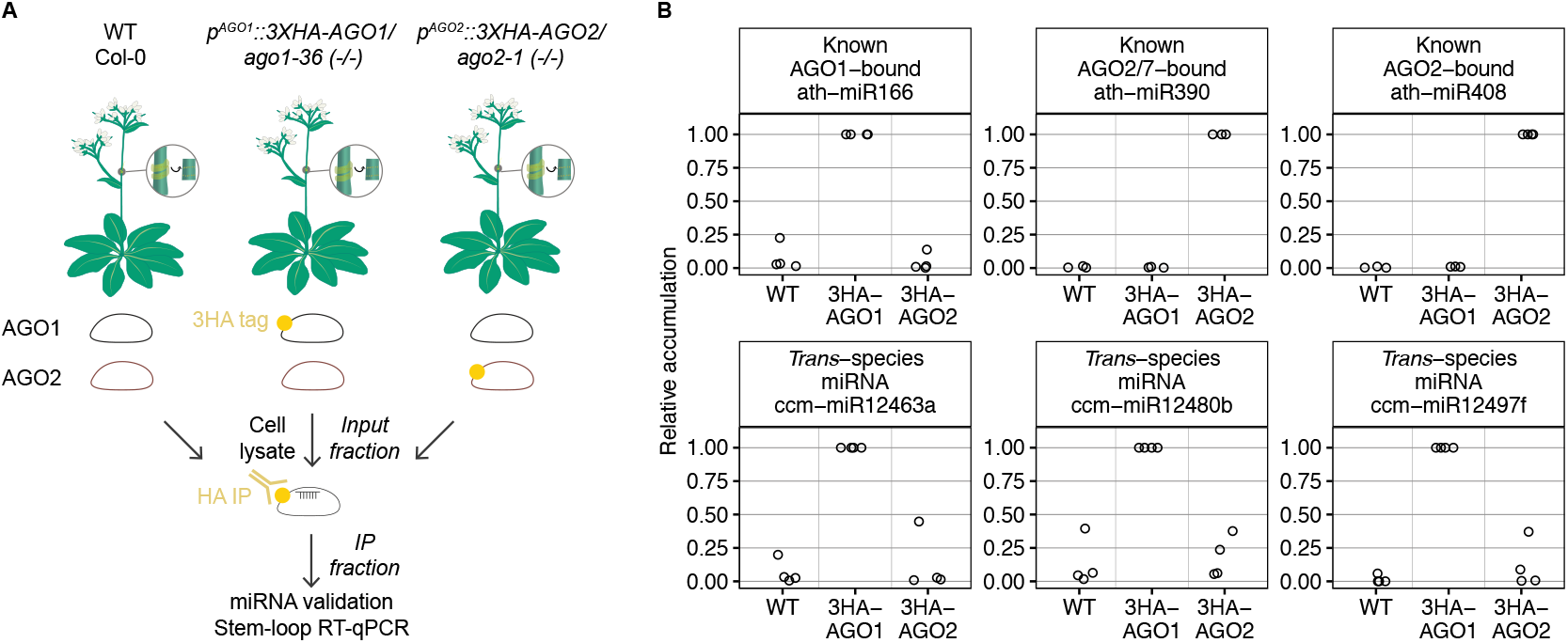
*C. campestris trans*-species miRNAs are preferentially loaded into host AGO1 over AGO2. **(A)** *C. campestris* was attached to three *A. thaliana* backgrounds: Col-0 (WT), 3XHA-tagged AGO1, and 3XHA-tagged AGO2. After ten days, miRNAs were quantified from total lysate (Input) and immunoprecipitated (IP) samples using RT-qPCR. **(B)** RT-qPCR result. Top: Control miRNAs are enriched only in their respective IPs, confirming specificity. Bottom: *Trans*-species miRNAs are strongly enriched in AGO1 IPs. Each dot represents a biological replicate consisting of ∼400 pooled *A. thaliana* stem segments beneath parasite attachment. Each condition included 3–4 biological replicates. Relative expression was calculated using the 2^−ΔΔCt^ (IP – Input fraction) method. 3HA-AGO1 served as the calibrator for ath-miR166 and all three *trans*-species miRNAs, while 3HA-AGO2 was used for ath-miR390 and ath-miR408. Data available in **Supplementary Table 1**.

## Discussion

Our working model of *trans*-species miRNA biogenesis and function in *C. campestris* is summarized in **Figure 6**. Transcription of *C. campestris trans*-species miRNA primary transcripts in haustoria precedes accumulation of mature miRNAs by 24-48 hours. Primary transcripts are processed by *C. campestris* DCL1. The resulting *trans*-species miRNAs avoid AGO loading inside of *C. campestris* and instead remain in miRNA/miRNA* duplex form until export. Upon arriving in host cells, they are loaded onto host AGO1 where they function to direct mRNA slicing and secondary siRNA biogenesis of targets. This mechanism allows *C. campestris* to produces a class of exclusively mobile miRNAs that are somehow pre-programmed for export, avoiding self-targeting, and only gaining function once inside the host.

**Figure 6.**
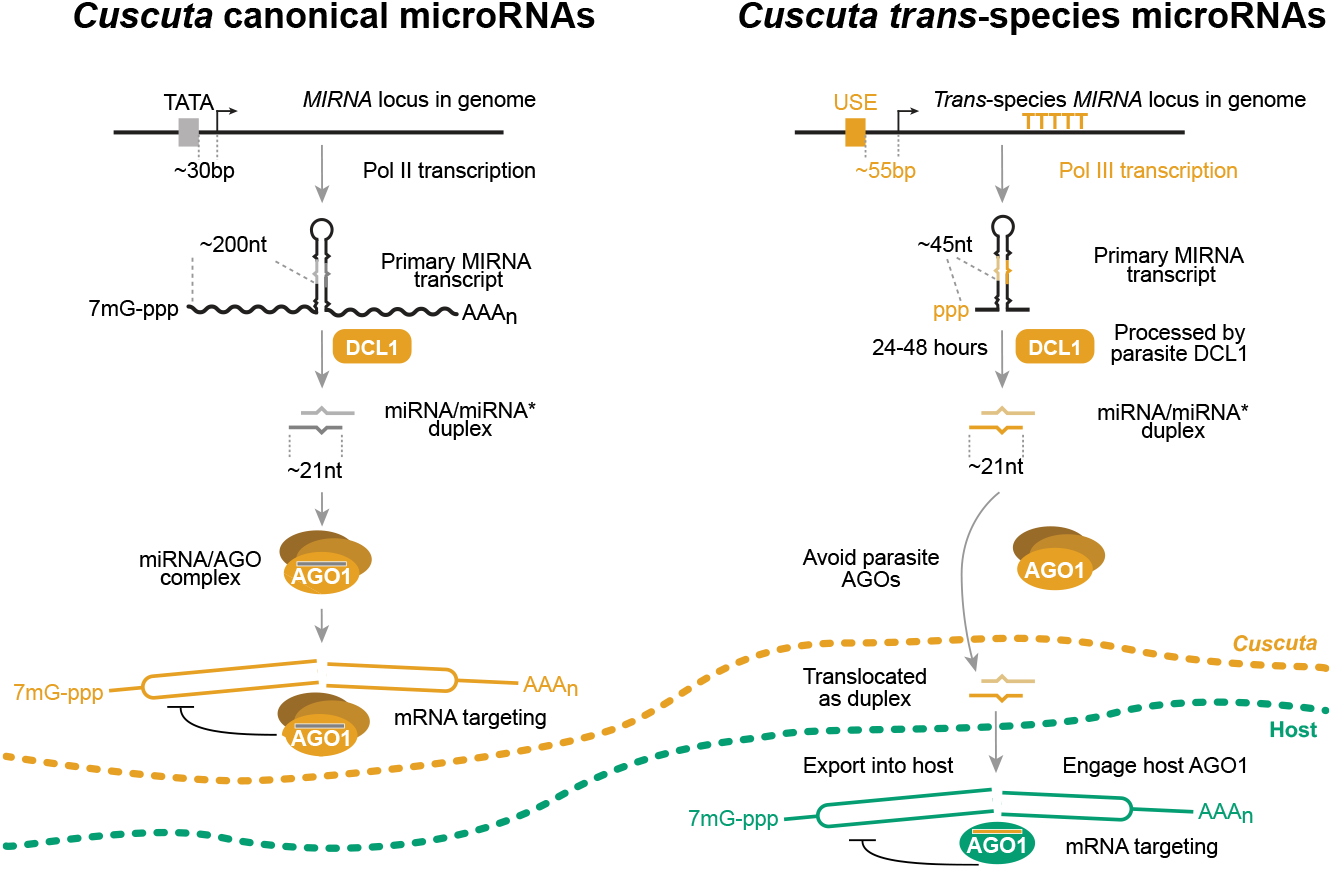
Biogenesis, export, and function of *C. campestris trans*-species miRNAs. Schematic summarizing our current model of miRNA biogenesis, export, and functions in the parasitic plant *C. campestris. Trans*-species miRNAs are produced by parasite DCL1, selectively avoid loading onto parasite AGO proteins, remain as duplexes until export, and only become functional in the host using host AGO1.

*Trans*-species miRNAs accumulate most strongly in haustoria but are also present at ∼10-fold lower levels in adjacent host tissues, spreading in both directions from the haustorium (**Fig. 1**). Their accumulation drops sharply at the basal petiole, suggesting limited long-distance mobility in the host (**Fig. 1**). Detection of *trans*-species miRNAs in adjacent host tissues allowed study after export, complementing host-free *in vitro* haustoria where miRNAs exist before export. Both systems provided single organism contexts for investigation: *in vitro* haustoria without host, and host tissue without parasite.

Primary transcripts peak 24–48 hours before their corresponding mature miRNAs and accumulate most strongly in haustoria, matching the highest mature miRNA levels (**Fig. 2C, D**). This suggests transcriptional regulation in both temporal and spatial dimensions. However, miRNA accumulation also reflects processing, AGO loading, turnover, and decay rates. Thus, higher mature miRNA abundance could arise from stronger transcription, more efficient dicing, or reduced turnover of transcripts that are constitutively produced at a basal level. Indeed, primary transcripts remain detectable across stages and tissues, including seedlings and shoot tips where mature miRNAs are absent (**Fig. 2C, D**), suggesting ongoing basal transcription. The upstream sequence element (USE) identified in most *trans*-species *MIRNA* loci is very similar to the constitutive U6 snRNA promoter^18^, consistent with a model where loci are transcriptionally “on” by default, while downstream regulatory layers refine mature miRNA accumulation through processing or degradation. Biologically, this default ‘on’ mode may benefit *C. campestris* by maintaining a basal pool of precursors that primes haustoria-competent tissues—such as seedlings and shoot tips—for rapid maturation of miRNAs once host contact initiates haustorial development. In *A. thaliana* stems near the attachment, mature miRNAs are present without detectable primary transcripts (**Fig. 2D**), supporting parasite origin and direct transfer rather than host processing.

*C. campestris trans*-species miRNAs are produced by parasite DCL1 but only become AGO bound after their arrival in the host. The three representative miRNAs that we examined were all strongly enriched in host AGO1 immunoprecipitates, but not in host AGO2 immunoprecipitates (**Fig. 5B**). However, we cannot at this point conclude that host AGO1 is the sole host AGO of importance. The three miRNAs examined all have a 5’-U, a feature that is highly preferred by AGO1. 5’-U miRNAs are the majority of confirmed *C. campestris trans*-species miRNAs, but a substantial minority of them have a 5’-C^18^. *A. thaliana* AGO5 primarily binds 5’-C RNAs and associates with both viral-derived siRNAs and some miRNAs^28–30^. Thus, we consider host AGO5 as another possible candidate for binding of some *C. campestris trans*-species miRNAs.

The difference between canonical miRNAs and *trans-*species miRNAs was striking: Within host-free *in vitro* haustoria, canonical *C. campestris* miRNAs are AGO-bound, while *trans-*species miRNAs are not (**Fig. 4E**). What accounts for this selective avoidance of AGO-loading by *C. campestris trans-*species miRNAs? Covalent nucleotide modifications are one possibility. Pseudouridine in miRNAs and siRNAs is associated with mobility^31^, while 5-methylcytosine modifications within mRNAs correlate with mRNA transport across graft junctions^32^. Similarly, tRNAs are heavily modified, and both full-length tRNAs and tRNA-derived fragments can move systemically through the phloem^33^. Addition of tRNA-like structures to non-mobile mRNAs can increase their systemic transport across grafting junction^34^ as well as into *C. campestris* from hosts^35^. The *Cuscuta*-host vasculature resembles a natural graft, and mechanisms that facilitate RNA transfer across graft junctions may also support *trans*-species movement. *C. campestris trans*-species miRNAs are transcribed from Pol III promoters^18^, which are also used by tRNAs. Future studies should prioritize investigation of RNA modifications on *trans-*species miRNAs.

These observations invite comparison with other cases of mobile small RNAs, both within a single plant and between plants and their associated organisms. Similar to the *C. campestris* system, *Botrytis cinerea* (gray mold fungus) sRNAs (Bc-sRNAs) are also processed by parasite DCLs, and after arriving in the host are bound to host AGO to effect gene silencing^6^. However, the *B. cinerea* AGO also binds mobile Bc-sRNAs and contributes directly to infection^36^. This seems to contrast with the avoidance of “self”-AGO loading we observed in *C. campestris*. Long and short distance mobility of siRNA-mediating gene silencing within a single plant has been tied to mobility of siRNA duplexes^37^. In this model, AGO proteins are not mobile, and once a given siRNA duplex is disassembled for AGO-loading, mobility ceases. This paradigm closely parallels our observations in *C. campestris* where *trans-*species miRNAs appear to accumulate as AGO-free miRNA/miRNA* duplexes within parasite tissues. Whatever the AGO-avoidance mechanism is, it may enable the *C. campestris trans*-species miRNA/miRNA* duplexes to easily enter the host.

*C. campestris trans*-species miRNAs largely avoid parasite AGOs and are exported as duplexes for selective loading into host AGO1. The biological benefit seems clear— avoiding self-targeting, which is particularly important given that parasitic plants and their plant hosts share greater sequence similarity than fungal pathogens and their host. A previous study^20^ demonstrated sequence dissimilarity between *Cuscuta* and host transcripts as one safeguard, but the complementarity scores are not zero, leaving some residual risk of self-targeting. Selective exclusion from parasite AGOs could therefore provide an additional layer of protection while at the same time encouraging movement into adjacent host tissues. The mechanism of selective AGO avoidance is a key unresolved question: it may involve differences between parasite and host AGO proteins, RNA modifications that enhance stability, or other undiscovered mechanisms. These questions define clear directions for future investigation.

## Online Methods

### Preparation of host plants and *Cuscuta campestris*

*N. benthamiana* and *A. thaliana* (Col-0) host plants were grown under long-day conditions (16 hours light / 8 hours dark) and watered weekly with a fertilizer supplement for 4 to 5 weeks prior to parasite attachment. *C. campestris* seeds were scarified in sulfuric acid for 1 hour, with gentle swirling every 20 minutes to ensure even exposure. After scarification, seeds were placed on moist tissue paper folded into a Petri dish and incubated at 30°C under long-day conditions supplemented with far-red light for 2.5 to 3 days. For host-induced haustoria, seedlings were taped to the host stem or petiole and incubated under far-red light enriched conditions for at least a week, until haustoria were firmly formed. *In vitro* haustoria were induced as previously described^38^. Briefly, *C. campestris* seedlings were sandwiched between 3% agar gel and 6 microscope slides. The sandwiches were exposed to far-red light enriched environment for 1 hour then immediately transferred to continuous blue light. Induced haustoria were harvested after 96 hours.

### Construction and agroinfiltration of artificial miRNA vectors

Artificial microRNAs (amiRs) were designed using P-SAMS^39^. A combined transcriptome database was assembled from *C. campestris*^40^ and *N. benthamiana* LAB3.6^41^. Three initial amiR candidates specific for *CcDCL1* and another three specific for *NbDCL1* were initially selected. Each initial candidate was predicted to be specific for the intended *DCL1* and to have very poor complementarity to the other *DCL1*. The candidate amiR showing the strongest silencing effect in pilot experiments was used for downstream analyses. The control amiR was designed to target an irrelevant mRNA (the human erythropoietin receptor mRNA, NM_000121). Forward and reverse oligos were synthesized, annealed and cloned into the *AtMIR390a* backbone^42^. The recombinant vector was transformed into *Agrobacterium* strain GV3101 and infiltrated as previously described^43^. In brief, a single colony was used to inoculate LB broth and incubated overnight at 28°C with shaking at 250 rpm. Cells were collected by centrifugation (4,000 × g, 15 minutes), resuspended in infiltration buffer (MgCl, MES, acetosyringone, sterile water) and incubated in a cool, dark place for ≥ 4 hours before infiltration. *N. benthamiana* leaves were infiltrated using a 1 mL needleless syringe. Three leaves per plant were treated, specifically the first and second pairs of true leaves. Agroinfiltrated areas generally covered entire leaves; if complete coverage was not possible, the area near the leaf base was infiltrated to enhance systemic movement of amiRs^23^. Oligonucleotide sequences are listed in **Supplementary Table 2**.

### Analysis of miRNA:miRNA* ratios

Three biological replicate small RNA-seq datasets from *C. campestris* host-free *in vitro* haustoria^18^ (SRA accession numbers SRR19502459, SRR19502460, and SRR19502461) were retrieved as FASTQ files from the NCBI short read archive (SRA). Adapters were trimmed using ShortCut version 1.0 with default settings (https://github.com/Aez35/ShortCut). A non-redundant list of *C. campestris* mature miRNA and miRNA* sequences was curated based on the miRNA annotated^16^. Occurrences of each miRNA and miRNA* sequence in each trimmed FASTQ file were tallied and converted from raw reads to reads-per-million (RPM). The data were filtered to retain only miRNA families where the mature miRNA is empirically known through prior demonstration of active targeting. RPMs were tallied by family for mature miRNAs and miRNA*s separately; these tallies were used to calculate the percentage of miRNA for each family.

### Enrichment of AGO-bound miRNAs using TraPR

Host-induced haustoria were established on *A. thaliana* inflorescence stem for 10 days under far-red light enriched conditions. Each inflorescence stem supported approximately ten attachments. Stems were excised above and below each attachment site, avoiding inclusion of *C. campestris* tissue. Each stem segment measured approximately 2-3 millimeters in length. For each biological replicate, 30 to 40 stem segments (∼80 milligrams total) were pooled. For *in vitro* haustoria, a total of 60 to 70 *C. campestris* seedlings were used to induce approximately 50 milligrams of *in vitro* haustorial tissue for each biological replicate. Tissues were processed through TraPR columns following the manufacturer’s instructions (TraPR Small RNA Isolation Kit, Lexogen, #128.08) with minor modifications. Briefly, tissues were homogenized in 300 μL TLB buffer using a bead beater at high intensity for 90 seconds (host-induced haustoria) and 30 seconds (*in vitro* haustoria). Lysates were centrifuged at 10,000 × g for 5 minutes at 4°C and the supernatant was transferred to a clean low-protein-binding tube (Thermo, #88379). The beads were rinsed with 50 to 150 μL nuclease-free water, centrifuged again, and the supernatant was combined. This rinse-and-centrifugation step was repeated until at least 270 μL of clarified lysate was collected. A 20 μL aliquot was reserved as input fraction, and the remaining 250 μL was processed using columns and TraPR fraction was obtained. Total RNA was extracted from both the input and TraPR fractions, followed by RT-qPCR analysis.

### Immunoprecipitation of epitope-tagged AGOs

*P*^*AGO1*^*::3XHA-AGO1* / *ago1-36* (-/-)^44^ and *p*^*AGO2*^*::3XHA-AGO2* / *ago2-1* (-/-)^45^ plants were grown in long day conditions (16 hours light / 8 hours dark) until the inflorescence stems reached at least 15 centimeters. Three-day-old *C. campestris* seedlings were attached to the inflorescence stem, with 10 attachments per plant. *C. campestris* was allowed to establish and grow for 10 days. *C. campestris* tissue was then peeled away and the *A. thaliana* stem tissue beneath each attachment site was collected. Approximately 400 such stem segments were pooled for each biological replicate. Tissues were ground in liquid nitrogen with a mortar until a fine powder was obtained. The frozen powder was mixed with 1 mL 4°C IP buffer (10 mM Tris-HCl, pH7.5, 150 mM NaCl, 0.5 mM EDTA, 0.5% (v/v) Nonidet™ P40 Substitute, RNase Out (Invitrogen™, #10777019) and 1 tablet of cOmplete EDTA-free Protease Inhibitor Cocktail / 50 mL (Roche, #11873580001). Immunoprecipitation using HA-Trap Magnetic Agarose (ChromoTek, #atma) was then performed. Lysates were centrifuged at 10,000 × g, 4°C for 10 minutes and the supernatant passed through a cell-strainer (40 μm, Falcon, #352340). A 200 μL aliquot was reserved for the input fraction, and the remaining lysate was incubated with HA-Trap Magnetic Agarose while rotating end-over-end for 2 hours at 4°C. Beads were washed 5 to 6 times with 4°C IP buffer. To elute bound HA-AGO complexes, beads were incubated at 95°C for 5 minutes in nuclease-free water with a dry bath. Total RNA was then extracted from both the input and IP fractions, followed by RT-qPCR analysis.

### RNA extraction and RT-qPCR to quantify RNAs and miRNAs

Total RNA was extracted using the Quick-RNA Plant Kit (Zymo Research, #R2024), generally following the manufacturer’s protocol with minor adjustments based on downstream applications. For mRNA and miRNA detection, RNA was not subjected to DNAse I treatment. Primers for mRNA detection were designed to span an exon-intron junction, and genomic DNA contamination is not a concern for the stem-loop RT primers used for miRNA detection. For primary miRNA transcript detection, RNA was treated with DNase I (Zymo Research, #E1009-A) and further purified using the RNA Clean & Concentrator Kit (Zymo Research, #R1019). For AGO-bound RNAs (17-200 nucleotides), including miRNAs and U6 snRNA recovered from TraPR and IP experiments, RNA was concentrated with the RNA Clean & Concentrator Kits using the alternative protocol (addition of 1.5 volumes of ethanol prior to column binding) to enrich low-abundance AGO-associated RNAs.

Reverse transcription was performed using the LunaScript® RT Master Mix Kit (Primer-free, New England Biolabs, # E3025L). For mRNA detection, Random Primer Mix (New England Biolabs, # S1330S) was used as the RT primer. For primary miRNA transcripts, gene-specific RT primers were used due to lack of poly-A tails. RT reactions followed the manufacturer’s protocol (2 minutes at 25°C, 10 minutes at 55°C, 1 minute at 95°C and then held at 4°C). For mature miRNA detection, primer design and RT conditions followed the previously described protocols^46,47^ with minor modifications: 20 minutes at 16°C, 60 minutes at 42°C, 5 minutes at 80°C and then held at 4°C. qPCR was performed using Luna Universal qPCR Master Mix (New England Biolabs, #M3003). For mRNAs and primary miRNA transcripts, the program followed manufacturer’s protocol (60 seconds at 95°C, followed by 40 cycles of 15 seconds at 95°C, 30 seconds at 60°C, with a standard melting curve from 65°C to 95°C). For mature miRNA, qPCR was run with 30 seconds at 95°C, followed by 40 cycles of 5 seconds at 95°C, 15 seconds at 52°C or 55°C (depending on the Tm of the universal reverse primer), and 10 seconds at 70°C, with standard melting curve from 65°C to 95°C. Oligonucleotide sequences are listed in **Supplementary Table 2**.

## Author Contributions

YCN performed most experiments, prepared all figures, and drafted the manuscript. MJA analyzed miRNA / miRNA* ratios and edited the manuscript. Both YCN and MJA jointly developed the experimental plan.

## Acknowledgements

We thank Xuemei Chen (Peking University) for the gift of *p*^*AGO1*^*::3XHA-AGO1* / *ago1-36* seed, and James Carrington (Donald Danforth Plant Science Center) for the gift of *p*^*AGO2*^*::3XHA-AGO2* / *ago2-1* seed. We also thank Teh-Hui Kao (Pennsylvania State University) and Linhan Sun (Pennsylvania State University) for providing guidance and equipment / reagents for the immunoprecipitation experiment. This work was supported by an award from the US National Science Foundation (Award #2003315) to MJA.

## Ethics Declarations

The authors declare no competing interests

## Supplementary Tables

**Supplementary Table 1**. Plotted data underlying all figures (Excel format).

**Supplementary Table 2**. Oligonucleotide sequences (Excel format).

## Extended Data Figures

**Extended Data Figure 1.**
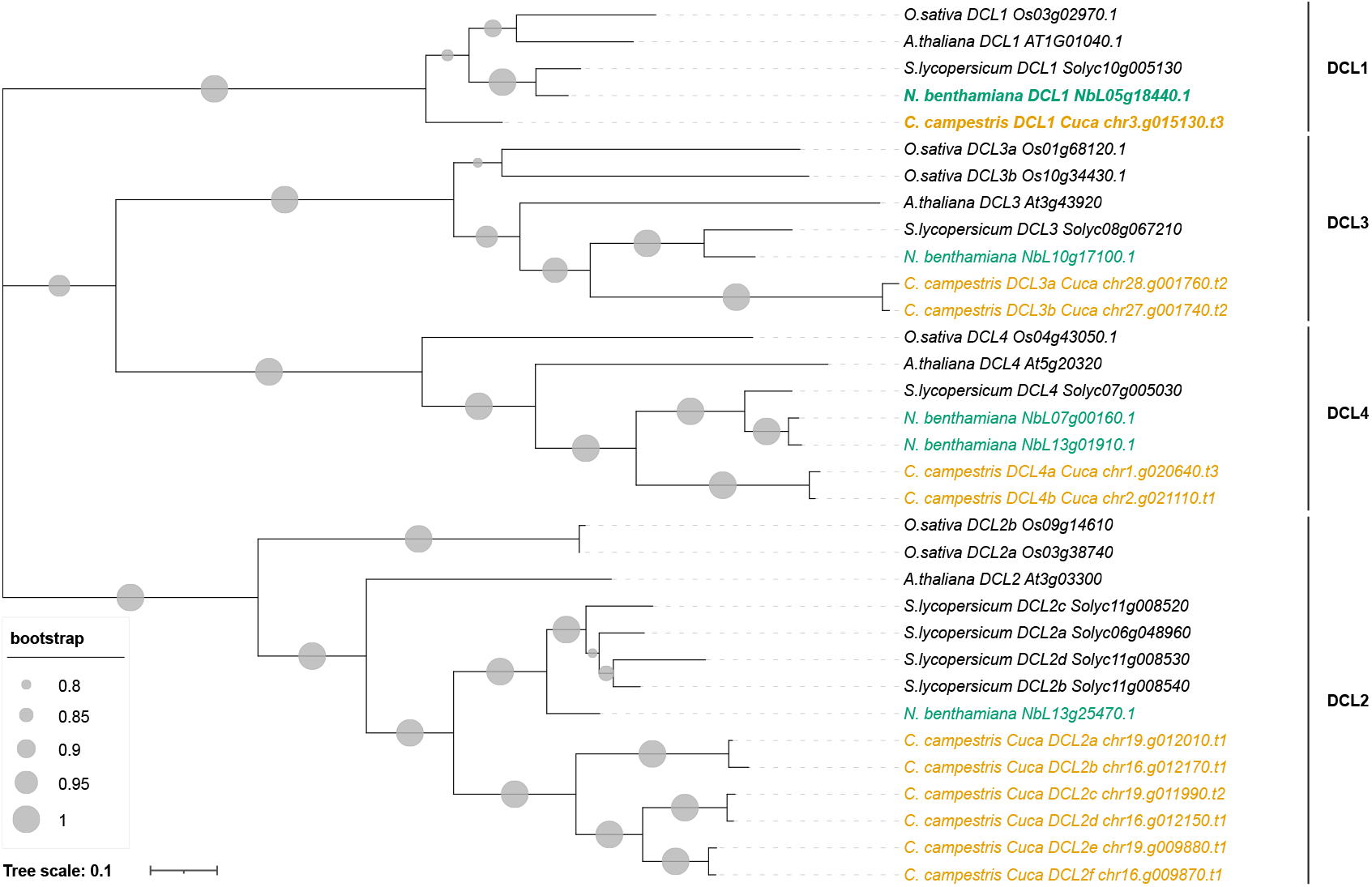
Phylogenetic analysis of DICER-LIKE (DCL) proteins in *C. campestris*. DCL protein sequences from *Oryza sativa*^48^, *Arabidopsis thaliana*^49^, *Solanum lycopersicum*^50^, *Nicotiana benthamiana*^41^, and *Cuscuta campestris*^16^. DCLs from *C. campestris* and *N. benthamiana* are highlighted with yellow and green, respectively. The silencing targets, *CcDCL1* and *NbDCL1*, are bolded and occur as single copies. The evolutionary history was inferred with the Maximum Likelihood method and Jones-Taylor-Thornton model. The tree is drawn with branch lengths measured in the number of substitutions per site. The bootstrap support values represent 500 replicates.

**Extended Data Figure 2.**
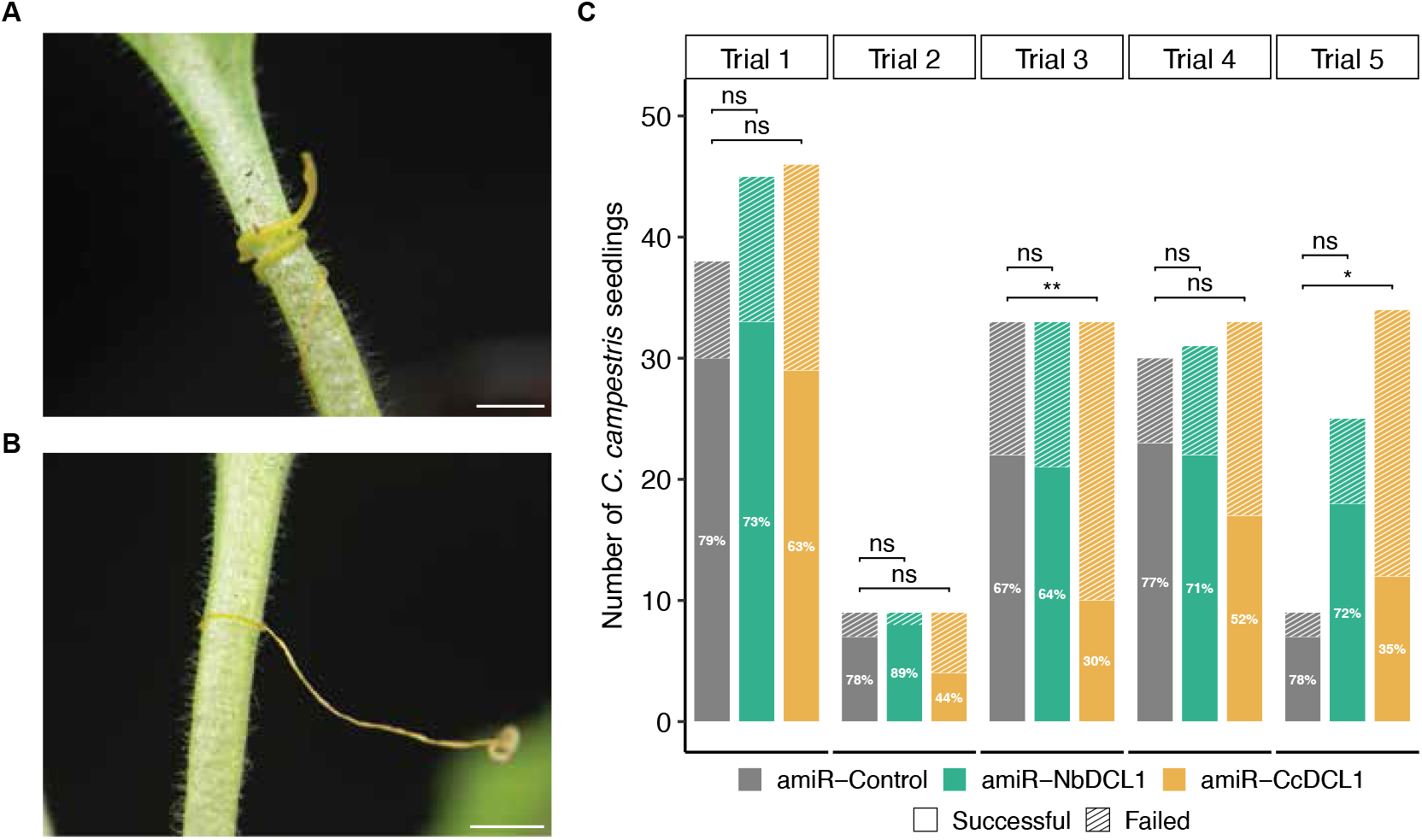
Reduced *C. campestris* establishment on hosts expressing artificial miRNA against parasite *DCL1*. **(A)** Successful establishment observed after seven days of *C. campestris* seedling coiling. The haustorial interface tightly grips the *N. benthamiana* petiole, displaying swelling and new shoot emergence. **(B)** Failed establishment showing a shriveled *C. campestris* seedling, devoid of second-round coiling, swelling, or new shoot growth. **(C)** Quantification of successful (solid) and failed (stripped) establishment count from five trials. Pairwise comparisons between amiR-Control (gray) and amiR-NbDCL1 (green) or amiR-CcDCL1 (yellow) were performed using Fisher’s exact test. Lower establishment rate is observed when attaching *C. campestris* seedlings to *N. benthamiana* infiltrated with amiR-CcDCL1, compared to host infiltrated with amiR-Control or amiR-NbDCL1. Statistical significance is indicated: ** p < 0.01, * p < 0.05, ns not significant. Data available in **Supplementary Table 1**.

## Notes

### Competing Interest Statement

The authors have declared no competing interest.

